# Anisotropic Rod-Shaped Particles Influence Injectable Granular Hydrogel Properties and Cell Invasion

**DOI:** 10.1101/2021.09.23.461542

**Authors:** Taimoor H. Qazi, Jingyu Wu, Victoria G. Muir, Shoshana Weintraub, Sarah E. Gullbrand, Daeyeon Lee, David Issadore, Jason A. Burdick

## Abstract

Granular hydrogels have emerged as a new class of injectable and porous biomaterials that improve integration with host tissue when compared to solid hydrogels. Granular hydrogels are typically prepared using spherical particles and this study considers whether particle shape (i.e., isotropic spheres versus anisotropic rods) influences granular hydrogel properties and cellular invasion. Simulations predict that anisotropic rods influence pore shape and interconnectivity, as well as bead transport through granular assemblies. Photocrosslinkable norbornene-modified hyaluronic acid is used to produce spherical and rod-shaped particles using microfluidic droplet generators and formed into shear-thinning and self-healing granular hydrogels at low and high particle packing. Rod-shaped particles form granular hydrogels that have anisotropic and interconnected pores, with pore number and size, storage moduli, and extrusion forces influenced by particle shape and packing. Robust in vitro sprouting of endothelial cells from embedded cellular spheroids is observed with rod-shaped particles, including higher sprouting densities and sprout lengths when compared to hydrogels with spherical particles. Cellular invasion into granular hydrogels when injected subcutaneously in vivo is significantly greater with rod-shaped particles, whereas a gradient of cellularity is observed with spherical particles. Overall, this work demonstrates potentially superior functional properties of granular hydrogels with rod-shaped particles for tissue repair.

## Main Text

Injectable hydrogels in combination with therapeutic cells and growth factors that are delivered through minimally invasive techniques are emerging as functional options to enhance tissue repair.^[1,2]^ Such hydrogels have also been investigated as acellular matrices to support endogenous repair, an approach that relies on the body’s innate regenerative capacity and stem cell pool.^[3]^ For an endogenous repair strategy to succeed, host cells and blood vessels must invade and populate the injected hydrogel to create a bioactive microenvironment.^[4,5]^ However, traditionally used bulk hydrogels are often rigid and non-porous, which prevents host cells from invading.^[6]^ Strategies to overcome this include incorporating enzymatically or hydrolytically degradable sites within the hydrogel,^[7,8]^ fabricating hydrogels from decellularized extracellular matrices (dECMs) with inherent degradability,^[9]^ or combining with fast-degrading or dissolving porogens to introduce porosity over time.^[10]^ Despite improvements, degradation of synthetic hydrogels and the inclusion of poragens are associated with weakening of mechanical properties and may additionally compromise structural features that provide instructive physical signals to host cells, while batch inconsistencies and limited ability to manipulate properties affect utility of natural dECMs.

To overcome these limitations, granular hydrogels have been developed in recent years to allow decoupling of material degradability from microporosity. In addition, granular hydrogels present other desirable features including injectability, modularity, and ease of tuning physical properties.^[11]^ Granular hydrogels are assembled through the packing of smaller (10^1^-10^2^ µm) hydrogel microparticles into a cohesive unit that is held together by chemical or physical interactions. Additionally, granular hydrogels are inherently porous owing to the interstitial space that exists between neighboring particles, and this porosity allows significantly higher cellular invasion and wound resolution when compared to bulk non-porous gels.^[12,13]^ Thus, granular hydrogels are ideally suited to promote the invasion of cells and blood vessels when used as injectable scaffolds for tissue repair. These desirable characteristics have motivated the development and application of granular hydrogels towards various biomedical applications including drug delivery,^[14,15]^ immune modulation,^[13,16]^ modular fabrication of biomaterials including with 3D printing,^[17–19]^ and as cell-instructive scaffolds.^[20–22]^

While much of the recent work on granular hydrogels has focused on tuning particle surface chemistry,^[12]^ inter-particle crosslinking,^[23]^ and varying other physical properties including stiffness and degradability,^[24,25]^ the influence of particle shape on interstitial pore characteristics to guide cell invasion has remained relatively unexplored. In a recent study, we reported that polygonal particles created through bulk hydrogel fragmentation pack together to form granular hydrogels that have much smaller pores when compared to those made with spherical particles.^[26]^ Additionally, Kessel and colleagues have reported similar pore features in granular hydrogels composed of entangled micro-strands obtained by passing a bulk hydrogel through a porous mesh.^[27]^ While pore size can impact the rate and extent of cellular invasion,^[28]^ interconnectivity of pores in three dimensions to form paths of least resistance is essential to guide cell invasion and migration.^[29]^ Smaller pores (pore volume < particle volume) such as those observed with highly packed polygonal fragmented particles are typically closed-off in three dimensions, whereas larger pores (pore volume > particle volume) tend to be more interconnected.^[21,30]^ Optimizing pore size, shape, and interconnectivity are therefore crucial to support the invasion of cells and blood vessels within granular hydrogels.

We hypothesized that granular hydrogels made with precisely engineered anisotropic particles would provide the appropriate guidance cues and interconnected porosity to accelerate cell and vessel invasion. To overcome the laborious and iterative process of optimizing the fabrication and characterization of granular hydrogels made with a variety of particles experimentally, we turned to simulations to test our hypothesis and guide our material design. Specifically, we used the graphics and simulations software Blender to predict how changing particle shape from spheres to rods would influence the structural features of granular hydrogels such as pore interconnectivity (**Figure S1**). Monodisperse populations of non-deformable spheres and rods with matching cross-sectional diameters (1 unit), but varied aspect ratios (1 vs. 2.2) were randomly packed into simulated cubes (9 × 9 units) to mimic granular hydrogels that exhibited similar porosity (**Figure 1a, Figure S1**). To elucidate differences in pore interconnectivity, we simulated and tracked the movement of an array of small beads (diameter: 0.2 units) that were dropped onto the surface of granular hydrogels as they trickled down through the interstitial pores over a period of several hundred frames; the simulations were run until all beads had settled (**Figure S2**). Positional data gathered from these simulations revealed that beads reached significantly greater z-depths with rod-shaped particles when compared to spherical ones (**Figure 1b, Figure S2**). Further analysis revealed that although both types of granular assemblies had similar overall porosity, bead movement through packed spherical particles was severely and frequently hindered as evidenced by the number of times that representative beads stopped in their migratory tracks, potentially due to randomly oriented pores, higher particle (obstacle) density, or path tortuosity (**Figure 1b, Figure S3**). In comparison, beads moving through packed rod-shaped particles stopped fewer times and reached 100% z-depth in fewer frames (**Figure 1b**). This may be due to contact guidance cues, pore anisotropy and interconnectivity, and migratory paths of least resistance (**Figure S3)**. Collectively, these data suggest that compared to granular hydrogels made with spherical particles, those made with rod-shaped particles may have structural features including pore interconnectivity, anisotropy, and contact guidance cues that enhance bead transport in silico and would likely also support enhanced cellular invasion. Pore interconnectivity is considered a vital factor controlling cellular invasion within porous scaffolds,^[31]^ and these simulations could have important implications for the rational design of biomaterials that promote endogenous repair.

**Figure 1.**
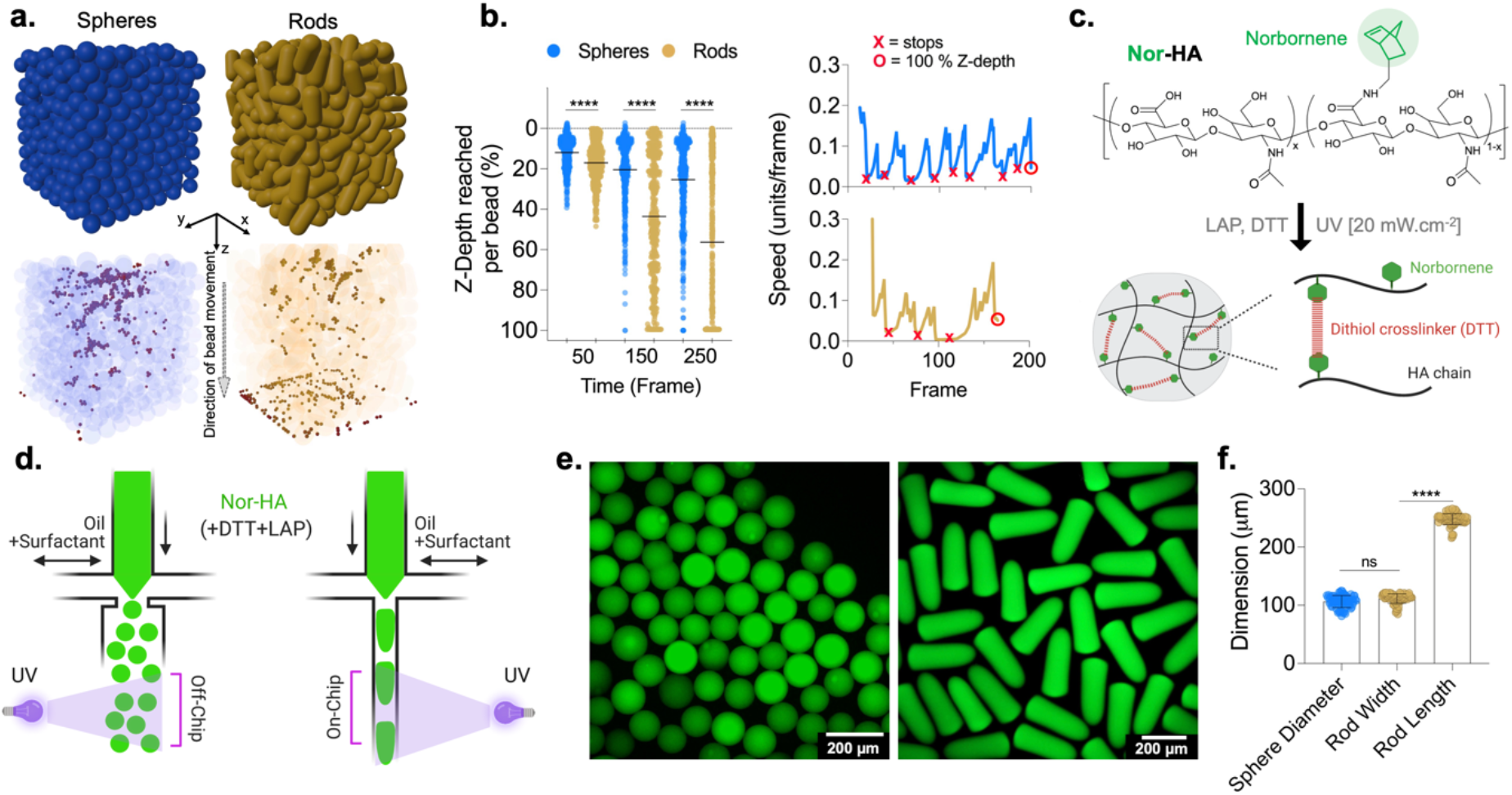
Varied particle anisotropy in granular hydrogels. (a) Simulations (Blender software) to characterize differences in pore interconnectivity in granular hydrogels made with spherical and rod-shaped particles, including the introduction of bead populations on the surface to investigate their transport through hydrogels. (b) Simulations predict improved interconnectivity with rod-shaped particles, based on depth of bead penetration (left) and travel speed (right). (c) Norbornene-modified hyaluronic acid (Nor-HA) structure and photocrosslinking via thiol-ene reaction. (d) Single-channel microfluidic droplet generators to fabricate Nor-HA particles of different shapes through either off-chip photocrosslinking (spheres) or on-chip confinement and photocrosslinking (rods). (e) Fluorescent images of monodisperse particle populations of spherical and rod-shaped particles that have (f) similar cross-sectional diameters (∼100 µm), but different aspect ratios (∼1 vs. ∼2.2; n=75). [****p<0.0001; ns = not significant]

Building on these simulations, we fabricated spherical and rod-shaped hydrogel microparticles to assemble and explore the influence of particle shape on granular hydrogel properties and cell invasion. PDMS-based single-channel microfluidic devices were used to create norbornene-modified hyaluronic acid (Nor-HA) microparticles. In these devices, surfactant-containing oil pinches off an aqueous stream of Nor-HA polymer to create spherical droplets, where the Nor-HA crosslinks through a thiol-ene reaction between norbornene functional groups on the HA backbone and dithiol crosslinkers in the presence of a photo-initiator and a UV light source (**Figure 1c**). Spherical particles were generated by exposing the droplets to UV off-chip (i.e., while droplets were flowing between the device outlet and collection vessel), whereas rod-shaped particles were formed by narrowing the width of the microfluidic channel downstream of the pinch-off point, causing the droplets to squeeze and adopt an elongated rod-like morphology while cured with UV on-chip (i.e., while rod-like droplets were flowing through the channel) (**Figure 1d**). This technique has previously been reported to make rods as well as flattened slab-shaped particles depending on the cross-sectional dimensions of the microfluidic channel.^[32]^ The microfluidic channel width governed the cross-sectional diameter of the rods, whereas the ratio of polymer to oil flow rate determined the diameter of the spherical particles (**Figure 1e**). Droplet deformation at the leading edge caused by differences in the viscoelasticity of the polymer and oil phase resulted in rods acquiring a bullet-like morphology, as previously reported.^[33]^ For this study, the experimental parameters were optimized to obtain spherical and rod-shaped particles with a matching cross-sectional diameter of ∼100 µm (**Figure 1f**). Rod-shaped particles had an aspect ratio of ∼2.2 compared to 1 for the spherical particles, and both groups of particles had a coefficient of variation of size <10%. While particle volume is not conserved between spheres and rods, here we focused on investigating the effect of changing particle shape by increase the aspect ratio from 1 to 2.2 while keeping the cross-sectional diameter consistent.

Granular hydrogels were assembled through centrifugation-mediated packing of spherical or rod-shaped particles (**Figure 2**). To visualize and quantify porosity, microparticles were resuspended in a solution of high molecular weight and fluorescent FITC-dextran prior to packing, and confocal microscopy was used to image interstitial pores. 3D rendering of the volumetric confocal stacks using IMARIS software revealed a network of interconnected pores within granular hydrogels that had distinct features depending on particle shape (**Figure 2a**,**b**). For instance, while particles in both groups were randomly packed, pores around spherical particles were seemingly randomly oriented, whereas those around rod-shaped particles were highly anisotropic and mimicked tubular features. The porosity of these volumetric reconstructions were obtained in IMARIS and validated against values obtained using a traditional method of thresholding individual 2D slices from confocal stacks (**Figure 2c, Figure S4**). One advantage of using centrifugation to assemble granular hydrogels from a suspension of soft and deformable particles is that centrifugation speed can be easily changed to control particle packing density. We exploited this to assemble granular hydrogels with either tightly packed or loosely packed particles, where porosity varied independent of particle shape and increased from ∼ 15% with tight packing to ∼ 25% with loose packing (**Figure 2b,c**). Interestingly, overall porosity remained unchanged based on particle shape, as long as the degree of packing (centrifugation speed) was consistent. In addition to overall porosity, the 2D thresholding of confocal slices allowed the quantification of the size and number of pores for each formulation. Comparison of these parameters from both groups showed that granular hydrogels fabricated with rod-shaped particles had a lower number of larger pores, whereas those made with spherical particles had a larger number of smaller pores (**Figure 2d-e**). Increasing the packing density also led to an increase in the number of pores but with significantly smaller average sizes. Together, these qualitative and quantitative data reveal stark differences in the structural features of interstitial pores that arise due to differences in the shape and packing density of particles within granular hydrogels.

**Figure 2.**
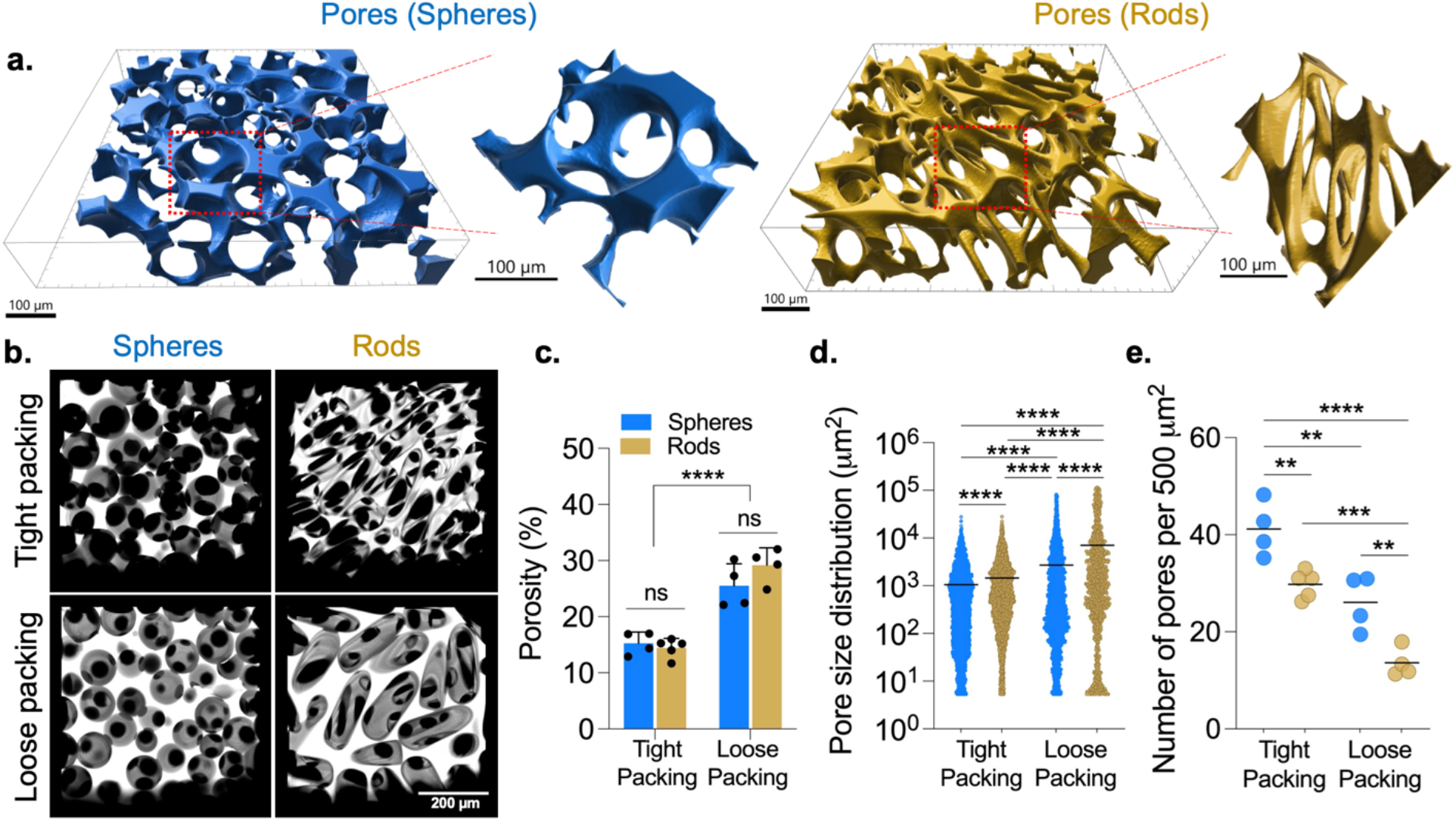
Particle shape impacts structural features of pores in granular hydrogels. (a) Visualization of interconnected pores within 3D volumetric regions of granular hydrogels made with spherical (blue) or rod (gold) shaped particles using confocal imaging and IMARIS software. (b) Changing particle shape and packing density (via centrifugation speeds) influence interstitial pores (bright = FITC dextran) that exist between neighboring microparticles (transparent), as observed in maximum projection images of confocal stacks. (c) Changing particle shape at similar packing density does not alter porosity, but changing packing density significantly modulates porosity from ∼15% (tight packing) to ∼25% (loose packing). 2D confocal slices are used to quantify the impact of particle shape and packing density on (d) pore size (cross-sectional pore area) and (e) pore number. Granular hydrogels with rod-shaped particles have a lower number of larger pores, whereas spherical particles result in a higher number of smaller pores. [**p<0.01; ***p<0.001; ****p<0.0001]

A major advantage of granular hydrogels for biomedical applications is their ease of injectability, which allows for placement into tissues and for 3D extrusion printing.^[17,18,27]^ Granular hydrogels show shear-thinning and self-healing behavior due to the displacement and flow of individual particles during extrusion through tight syringe needles or small catheters and the recovery of mechanical properties upon removal of strain. In a recent granular hydrogel study that compared the rheological behavior of spherical particles versus polygonal fragments, shape changes were found to significantly increase storage moduli and extrusion forces, independent of material composition and cross-linking mechanism, suggesting that particle shape and structural features directly influence hydrogel properties and injectability.^[26]^ Using oscillatory shear rheology, we found that all granular hydrogels developed in this study showed shear-thinning and self-healing behavior under periodically applied regimes of high (500 %) and low (1 %) strains, independent of particle shape or packing (**Fig. 3a**). Moreover, all granular hydrogels tested showed a complete recovery of their mechanical properties upon removal of high strain. Granular hydrogels behave like solids under low strains and the storage modulus (G’) indicates their resistance to mechanical deformation. While packing density affected storage moduli for both particle shapes (G’_tight packing_ > G’_loose packing_), granular hydrogels made with rod-shaped particles had significantly higher storage moduli when compared to those made with spherical particles at the same packing density, likely due to differences in deformation, sliding, or interlocking of particles under the application of shear (**Fig. 3b**). At high strains, granular hydrogels undergo yielding, which allows them to flow (i.e. G”>G’) through narrow openings such as syringe needles (**Fig. 3c**). Interestingly, the yield strain values for granular hydrogels made with rod-shaped particles were lower (tight packing) or similar (loose packing) to those made with spherical particles (**Fig. 3d**)

**Figure 3.**
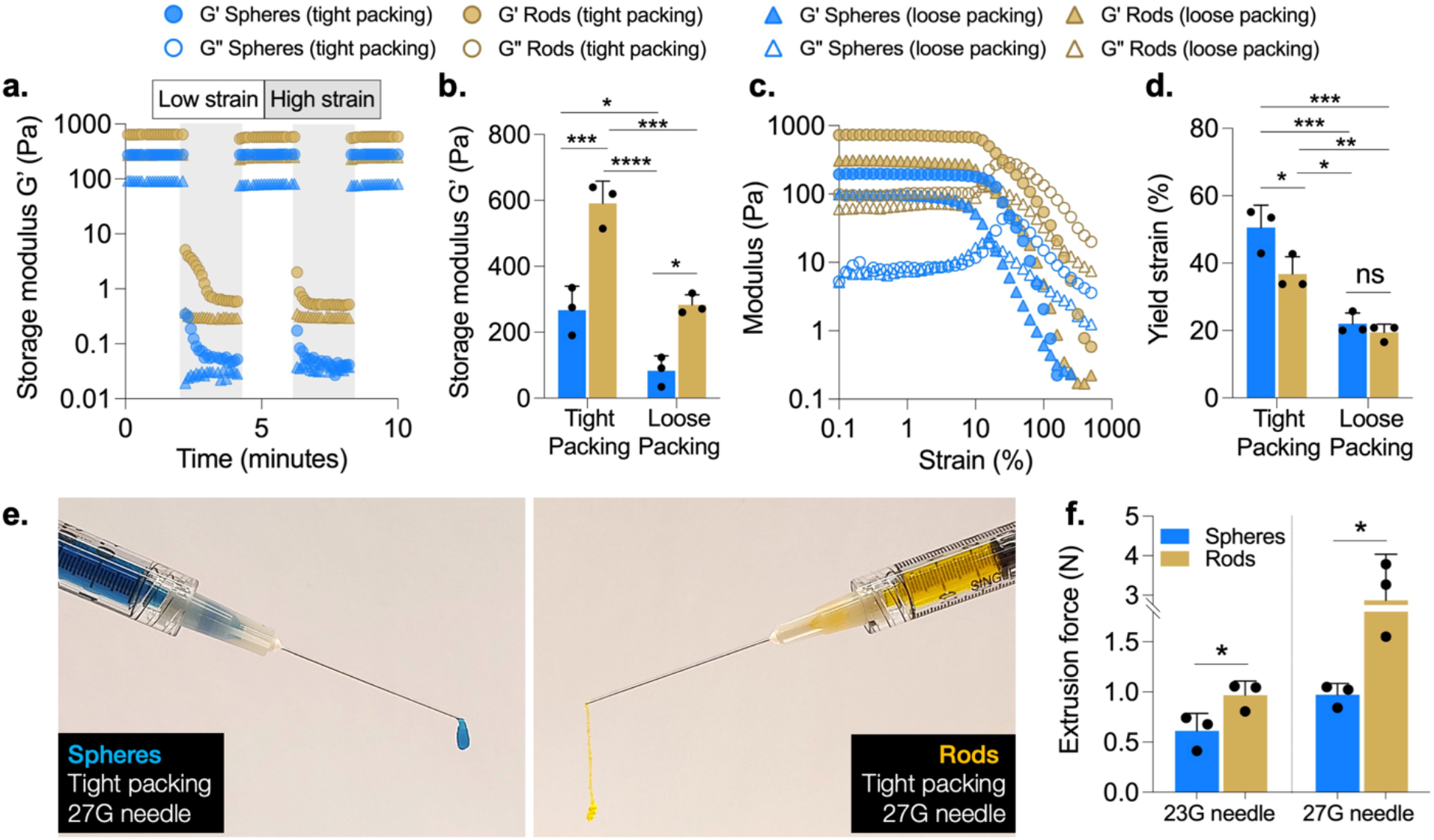
Particle shape impacts rheological behavior and injectability of granular hydrogels. (a) Representative time sweeps with periodically applied regimes of high (500 %) and low (1 %) strains showing shear-thinning and self-healing of granular hydrogels. (b) Storage moduli (G’; derived at 1 % strain) of granular hydrogels are significantly greater with rod-shaped particles (tight packing: ∼590 Pa; loose packing: ∼280 Pa) compared to spherical ones (tight packing: ∼270 Pa; loose packing: ∼80 Pa). (c) Representative strain sweeps (0.1 % to 500 %) showing shear-yielding properties of granular hydrogels with yield strains (% strain at G”>G’) ranging between 10 and 100 %. (d) Yield strains vary significantly with changing packing density while remaining comparable for different shapes at same packing density (tight packing: ∼50 % rods vs. ∼37% spheres; loose packing: ∼22 % rods vs. ∼20 % spheres). (e) Images showing extrusion of granular hydrogels through clinically relevant 27G syringe needles. (f) Extrusion forces vary with particle shape and needle diameter; rod-shaped particles and smaller needles (27G) require higher forces than spherical particles and larger needles (23G). [*p<0.05; **p<0.01; ***p<0.001; ****p<0.0001]

Differences in the storage moduli of rod-based and sphere-based granular hydrogels also determined their behavior when extruded through surgically relevant 27G needles (**Fig. 3e**). While granular hydrogels composed of spherical particles behaved like a highly viscous material upon extrusion, those made of rod-shaped particles extruded as continuous strands. Additionally, extrusion forces were dependent on particle shape (forces were significantly higher for rod-based samples) and needle size (forces were higher for smaller needle diameters). Taken together, this analysis of the rheological behavior of granular hydrogels made with spherical and rod-shaped particles demonstrate their general ease of injectability, the tunability of their bulk mechanics under shear, and differential properties based on particle shape and packing. These unique rheological properties make them suitable for a broad range of biomedical applications, particularly where their minimally invasive delivery could confer a substantial clinical advantage.

In contrast to traditionally used non-porous bulk hydrogels, highly porous granular hydrogels support the invasion of cells and blood vessels within their volume, thereby fostering the requisite biological activity to promote endogenous tissue repair.^[4,14,21]^ While several previous studies have investigated the adhesion, spreading, and migration of single cells throughout granular hydrogels as a function of particle properties, it remains unknown how endothelial cell sprouting – a process that mimics angiogenesis – is influenced by granular hydrogel properties.^[34,35]^ This investigation is challenging in vitro because cells preferentially adhere and proliferate on particle surfaces, which reduces their ability to form lumen-like sprouts. To overcome this, we established a 3D in vitro endothelial sprouting assay where multicellular spheroids composed of endothelial and mesenchymal cells are produced and suspended within an interstitial matrix that is embedded within centrifugally packed granular hydrogels (**Figure 4a** and **Figure S5**). Angiogenic factors introduced into culture media stimulate the formation of endothelial sprouts that then extend into the interstitial volume between the packed particles, with particle packing and interstitial volume unaffected by the presence of the interstitial matrix (**Figure S6**). While this setup does not mimic the complex biological conditions such as matrix deposition on implanted materials that may modulate angiogenesis in vivo, this assay can be used to screen how granular hydrogel features such as porosity and interconnectivity affect endothelial sprouting behavior.

**Figure 4.**
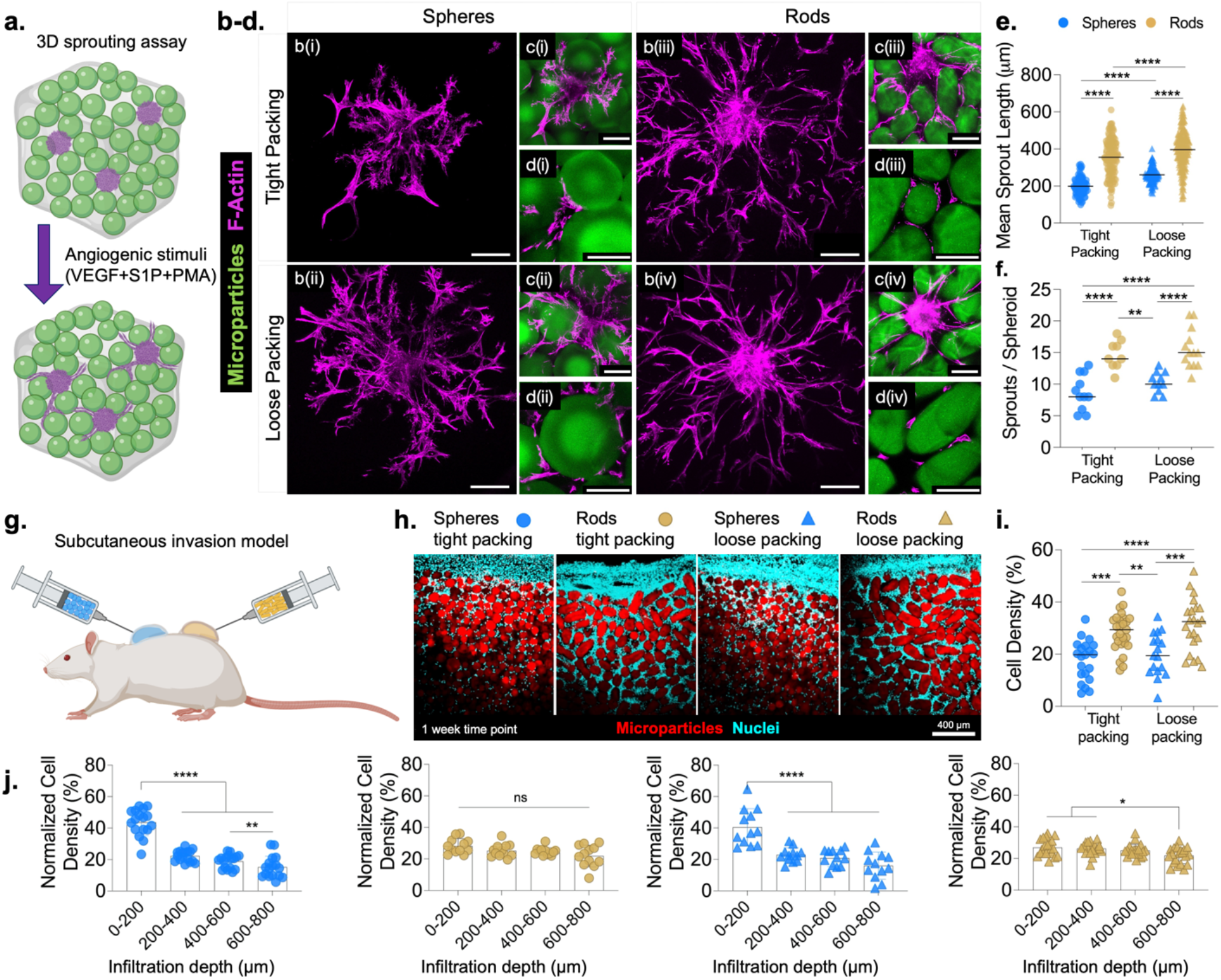
Particle shape impacts in vitro endothelial sprouting behavior and in vivo cellular invasion. (a) Schematic of the in vitro 3D sprouting assay where spheroids (magenta) are embedded within an interstitial matrix (grey) and distributed throughout packed granular hydrogels made of either spherical or rod-shaped particles (green); sprouts emerge in response to angiogenic stimuli and are imaged and analyzed at day 3. (b-d) Cells sprouting from spheroids of endothelial and mesenchymal cells and extending through available interstitial space within tightly packed (i, iii) or loosely packed (ii, iv) granular hydrogels made of spherical (i, ii) or rod-shaped (iii, iv) microparticles. Representative maximum projection images showing (b) F-Actin stained sprouts extending from spheroids; (c) radial sprouts invading interstitial pores surrounding the spheroid; (d) clearly visible lumens in sprouts extending through interstitial pores. Granular hydrogels with rod-shaped particles facilitate significantly greater (e) sprout lengths, and (f) sprout density per spheroid. (g) Endogenous cell invasion into granular hydrogels is assessed in a subcutaneous model. (h,i) One week after injection, significantly higher cell density is observed with rod particles compared to spherical ones in both tightly packed (25% in rods vs. 15% in spheres) and loosely packed (25% in rods vs. 15% in spheres) granular hydrogels. (j) Gradient in cell invasion is observed with spherical particles where cell density is highest in the top 200 µm and significantly declines thereafter; with rod particles, no gradient exists with uniform cell density up to a depth of 800 µm. [*p<0.05; **p<0.01; ***p<0.001; ****p<0.0001]

We used this assay to investigate how particle shape and packing density influence endothelial cell sprouting behavior, including sprout length and sprout density (i.e., number of sprouts per spheroid). Because sprouts extend outwards from the spheroid, other endothelial network features that are typically observed in 2D tube formation experiments including branching points and junctions were not observed with this assay. Our data show that both particle shape and packing density significantly affect sprouting behavior (**Figure 4b-f**). Sprout length and density were higher with rod-based granular hydrogels when compared to those containing spherical particles at both packing conditions tested. These data point to a likely critical role of the contact guidance cues provided by the major axis of the rod-shaped particles and the anisotropic pores that were identified earlier (**Figure 2a**) in supporting the directional growth of endothelial sprouts (**Figure 4c iii-iv**). In contrast, the smaller sprout lengths observed with sphere-based granular hydrogels likely reflect the randomness of pore orientation, as well as the comparatively more tortuous paths that connect those pores accentuated by the surface curvature of the spherical particles, as determined earlier through simulation and experiments (**Figure S3, Figure 2**). As a result, sprouts that are blocked by the presence of spherical particles in their path have to migrate around the particle until an interstitial pore opening is reached (**Figure 4d i-ii**). As expected, packing density also modulated sprouting behavior with smaller sprout lengths observed with tight packing, likely due to the effects of particle packing on interstitial porosity.

Ultimately, we envision a use for these granular hydrogels as injectable acellular materials that facilitate endogenous tissue repair through the recruitment of cells into interstitial pores. To demonstrate endogenous cell recruitment in vivo, we injected acellular granular hydrogels fabricated with either spherical or rod-shaped particles and packed at two densities into subcutaneous pockets of adult athymic rats, and histologically analyzed cell invasion at one week after injection (**Figure 4g**). These granular hydrogels were prepared without the interstitial matrix that was used for in vitro studies. Importantly, these particles were fabricated with DTT as the dithiol crosslinker, allowing us to investigate cell invasion into non-degradable granular hydrogels and we confirmed that the needle injection process does not affect the measured porosity of granular hydrogels (**Figure S7**). After one week, we observed significantly higher infiltrated cell densities into rod-based granular hydrogels when compared to spherical ones (**Figure 4h-i**). However, differences in particle packing did not lead to observable differences in cell invasion when using the same particle shape, further demonstrating the importance of changing particle shape and anisotropic cues in guiding a strong cellular invasion response. Beyond quantifying overall cell density, sample cryosections were assessed for cellularity at regions of interest each measuring ∼200 µm in depth from the material-tissue interface. With spherical particles, a clear gradient was observed with the highest cell density fraction recorded for the first 200 µm from the interface, which significantly decreased thereafter (**Figure 4j**). In contrast, no gradient was observed with rod particles, particularly up to 600 µm from the material-tissue interface, suggesting an uninhibited invasion of cells into rod-based granular hydrogels. Taken together, the in vitro and in vivo findings described in this section demonstrate significantly greater and uninhibited invasion of endogenous cells into granular hydrogels fabricated with rod-shaped particles highlighting the impact of particle shape in a biological context.

In conclusion, this paper describes the influence of changing particle shape from isotropic spheres to anisotropic rods on the porosity and injectability of granular hydrogels and their ability to support endothelial sprouting and cellular invasion. Simulations with Blender software provided in silico evidence that pore interconnectivity and bead transport through granular assemblies are improved with rod-shaped particles. Experimentally, changing particle shape from spheres to rods impacted several structural characteristics of granular hydrogels including pore shape, size, and number, and introduces directional contact guidance cues. Granular hydrogels made with rods have higher storage moduli and require higher extrusion forces but remain injectable through syringe needles without compromising on porosity. Furthermore, rod-shaped particles supported robust endothelial sprouting in vitro compared to spheres, with both particle shape and packing influencing sprout lengths and sprouting densities. Lastly, acellular granular hydrogels made with rod-shaped particles supported significantly greater and uninhibited invasion of endogenous cells when injected subcutaneously. Altogether, these results point to a potentially superior class of functional granular hydrogels made with anisotropic rod-shaped particles.

Future studies can build on the findings reported here by exploring how other parameters closely related to particle anisotropy influence granular hydrogel properties and function. For example, it is unknown whether aspect ratio of rod-shaped particles or diversity in shapes of anisotropic particles modulates porosity and cell-guidance cues while retaining ease of injectability. To answer this question, particle fabrication techniques will need to be improved such that they allow the high-throughput creation of particles with consistent material compositions, mechanics, and dimensions with the ability to tune these parameters as desired. Besides the method used in this paper, a number of other approaches have also been reported to produce anisotropic hydrogel microparticles. For example, Krueger et al. used compartmentalized jet-polymerization on microfluidic chips to produce a stream of anisotropic particles with varying aspect ratios,^[36]^ whereas Le et al. utilized photo-lithography to create soft and degradable rods.^[37]^ While these techniques result in similar outcomes in terms of particle anisotropy, properties, and downstream applications, their scalability for high throughput production is lacking and future work in this area will rely on innovations in processing techniques to produce the high particle numbers required for scale-up to large animal and eventually human studies.^[38]^ The iterative and laborious nature of device fabrication and microparticle production can be circumvented to some degree by harnessing the power of in silico studies.

Although simulations provided a necessary impetus to the work performed in this study, there are numerous opportunities to further optimize simulation conditions and enhance its predictive power, such as through creating deformable particles instead of rigid ones and through elaborating on the variations and heterogeneities observed with particle size and mixing of separate particle populations. Lastly, while the subcutaneous cell invasion model highlights important differences in the invasion of cells into granular hydrogels made with different shapes, a limitation is that it does not mimic a tissue injury environment. Future studies could therefore elaborate on the impact of particle shape and granular hydrogel properties on the numbers, identities, and kinetics of invading cells and establish how these relate to tissue repair outcomes. While we show that particle shape leads to differences in pore characteristics that impact cell invasion, other approaches to modulate porosity such as the mixing in of degradable particles could also be useful in creating space for cell invasion and expansion. We expect that high interest in the biomedical applications of granular hydrogels will lead to continued innovation in the design and applications of these functional materials for tissue repair applications.

## Experimental Methods

### Polymer synthesis

Norbornene-modified hyaluronic acid (Nor-HA) was prepared as previously described.^[38]^ Briefly, HA modified with tetrabutylammonium salt (HA-TBA) was dissolved in anhydrous dimethyl sulfoxide (DMSO). Dimethyl aminopyridine (DMAP), norbornene-2-carboxylic acid, and ditertbutyl dicarbonate (Boc_2_O) were added to the DMSO and allowed to react overnight at 45 °C. The reaction was quenched with a equal volume of cold water, and the Nor-HA product was then dialyzed for 14 days in DI water, frozen, lyophilized until dry, and stored at −20 °C until further use. To determine the degree of modification of the HA backbone with norbornene functional groups, lyophilized polymer were dissolved in deuterium oxide (D_2_O) at a concentration of 10 mg.mL^-1^ and analyzed using ^1^H NMR (Bruker NEO400).

### Simulations with Blender

Simulations were run in Blender 2.91 using the Bullet Physics engine (Figure S1a). Granular assemblies were simulated by randomly packing rigid body spheres (diameter = 1 unit) or rods (diameter = 1 unit; aspect ratio = 2.2) in a cubic container (volume = 729 units^3^) under the force of gravity using collision detection. Spherical collision shape was used for spheres; mesh collision shape was used for rods. Once settled, spheres or rods were joined into a single passive rigid body mesh object to form a granular assembly. Porosity of the granular assemblies was determined by using the Bool Tool to create 2D cross-sectional images and subsequently using a MATLAB code to calculate percent void space (white pixels, Figure S1b) in each image.

To investigate pore interconnectivity, spherical beads (diameter = 0.2 units) with active rigid body physics were dropped into the granular assemblies under the force of gravity using spherical collision shape. A total of 441 beads arranged in a plane were placed directly above the granular assemblies (z position = 0) and dropped under the force of gravity. In each simulation, 1000 frames were simulated such that all beads were settled. For each bead, object position data (xyz coordinates) was extracted at various time points. Once all beads were settled, percent z depth was determined for each bead (0% for bead z position = 0; 100% for bead z position = -9). Speed and tortuosity were also quantified using the extracted object position data.

### Granular hydrogels

Hydrogel microparticles were produced by flowing a mixture of Nor-HA (3% w/v), DTT (4 mM), and LAP (0.075% w/v) with or without 2 MDa FITC-Dextran in PBS through a PDMS-based single channel microfluidic device. The microfluidic device to produce spherical particles was fabricated as described before.^[17]^ Briefly, polydimethylsiloxane (PDMS; Sylgard 184, Dow Corning) was cast onto plastic molds that were fabricated via stereolithography (Microfine green; ProtoLabs) using a custom-made design. After overnight curing at 37 °C, the PDMS substrate was peeled off from the mold, punched with 1 mm diameter biopsy punches (Integra Miltex) to create inlet and outlet channels, cleaned, and plasma-bonded to a standard glass slide. Stainless steel dispensing needles (19G; McMaster Carr) were inserted into the inlet and outlet channels. Silicone tubing (Tygon; ABW00001, Saint-Gobain) was connected to the inlet and outlet needles prior to using the device. To fabricate microparticles, light mineral oil with 2 v/v % Span 80 was used as the continuous phase, and polymer precursor solution was used as the dispersed droplet phase. An oil flow rate of 40 μL.min^-1^ and a precursor solution flow rate of 5 μL.min^-1^ was used; this combination of flow rates had been optimized for the production of 100 µm diameter spherical particles. Droplets were exposed to 20 mW.cm^-2^ UV light (Omnicure S1500) near the collection tube (off-chip) to make spherical particles.

The microfluidic device to produce rod-shaped particles was fabricated in a similar way. Briefly, PDMS devices containing microchannels were fabricated following a standard soft lithography process: negative photoresist SU-8 2050 was spin coated on a silicon wafer to achieve a height that matched the width of the flow-focusing point (100 µm), followed by UV exposure using a mask aligner (SUSS Microtec MA6). After baking, the photoresist was developed in SU-8 developer to reveal the patterns, washed with isopropyl alcohol (IPA) and dried with Nitrogen gas. The hard master was vapor-coated with Trichloro(1H,1H,2H,2H-perfluorooctyl) silane (Sigma) in vacuum. PDMS was then cast onto the hard master, cured, peeled off, punched, and bonded following the same procedure described above. To fabricate rod-shaped microparticles, Nor-HA precursor solution was introduced to the device as the dispersed phase and emulsified by a continuous phase that was composed of perfluorinated oil HEF-7500 (3M) with 2 v/v % of the surfactant Krytox-157 FSH (Dupont). The flow rates of the prepolymer solution and oil were kept at 0.15 mL.h^-1^ and 1 mL.h^-1^, respectively. The portion of the microchannel downstream of the flow-focusing point was exposed to a 405 nm laser on a fluorescence microscope (on-chip) to solidify rod-shaped droplets. The microparticles were collected into HFE-7500 containing 20 vol.% of 1H,1H,2H,2H-Perfluoro-1-octanol (Sigma) to destabilize the surfactants. After removing the majority of the oil by centrifugation, the microparticles were washed sequentially with Hexane, 50:50 mixture of ethanol:DI water, and DI water to remove remaining oil, and transferred to PBS for further use. Between each step, a centrifugation at 5000 rcf for 3 minutes was performed.

The obtained particles were washed multiple times with PBS and 1% Tween-20 to remove any remaining oil and surfactant. Particles were then assembled into granular hydrogels by centrifuging particle suspensions at either 1000 rcf (loose packing) or 15000 rcf (tight packing) for five minutes. The supernatant was aspirated, and the granular hydrogel pellet was retrieved and handled further using a spatula.

### Characterization of granular hydrogels

To characterize porosity, particles without encapsulated FITC-dextran were resuspended in a 10 mg.mL^-1^ solution of high molecular weight FITC-dextran (2 MDa) in PBS before centrifugation-mediated packing. The granular hydrogel pellet was transferred to a sample holder that was prepared by punching out a cured PDMS slab and adhered to a standard glass slide. A confocal microscope (Leica TCS SP5) equipped with an Argon laser and a 25x water immersion objective lens was used to visualize the FITC-dextran present within interstitial pores. Multiple 3D stacks with average z-depths of ∼100 µm and inter-slice z-spacing of ∼5-7 µm were acquired at randomly chosen regions of interest. Microscopy files were further analyzed in FIJI software. The fluorescent signals in each slice within a stack were thresholded, smoothened, and the area fraction of the fluorescent signal was obtained using the in-built Analyze Particles function. The percentage of the fluorescent area fraction to the total area was calculated and termed porosity. Porosity values from successive slices were averaged to obtain overall porosity/interstitial volume. Additional outcomes from these analyses included total number and cross-sectional area of pores. IMARIS software package was used to visualize 3D porosity within granular hydrogels using the same confocal microscopy raw data sets. The surfaces feature was used with average feature size of 1.5 µm to recreate volumetric pores.

To characterize rheological behavior, granular hydrogels that were packed through centrifugation were transferred to an oscillatory shear rheometer (AR2000, TA Instruments) equipped with a 20 mm parallel plate geometry set at a 1200 µm gap. All tests were conducted at room temperature. Strain sweeps (1−500% strain, 1 Hz) were used to assess shear yielding properties. For flow characterization, viscosity was measured with a continuously ramped shear rate (from 0 to 50 s^−1^). For shear recovery experiments, low (1%) and high (500%) strains were periodically applied at a frequency of 1 Hz.

To quantify extrusion forces, granular hydrogels that were packed through centrifugation were transferred to a 1 mL Luer Lock syringe equipped with either a 23G or 27G needle. The syringes were loaded onto a syringe pump, and a round force-sensitive resistor (Interlink 402) was placed in between the syringe plunger and the pump. Data was acquired using a custom setup built with an Arduino Uno Rev3. The pump was switched on with the extrusion rate set to 10 mL h^-1^. Voltage output signal was recorded from the Arduino IDE serial monitor. Force calibration curves were generated earlier using standard weights. Extrusion forces recorded with the different granular hydrogel groups were converted using this calibration curve, and the last 30 seconds of a 1 minute total measurement period was used to calculate the mean extrusion force.

### In vitro sprouting assay

Multicellular spheroids consisting of human umbilical vein endothelial cells (HUVECs; Lonza) and human mesenchymal stromal cells (MSCs) in a 2:1 ratio were prepared by seeding cells on AggreWell™ 400 templated agarose wells and culturing overnight. Cell concentrations were adjusted to target ∼1000 cells per spheroid. Spheroids were washed twice with PBS prior to use. An interstitial matrix precursor solution was produced by combining 1.8% w/v Nor-HA, 0.5 mM thiolated RGD (GCGYGRGDSPG, Genscript), 3.5 mM thiolated MMP-degradable peptide crosslinker (GCNSVPMSMRGGSNCG, Genscript), and 0.05% w/v LAP photoinitiator) in PBS. Hydrogel microparticles (with encapsulated high molecular weight FITC-Dextran) were re-suspended in this precursor solution and rigorously mixed on an orbital shaker for 20 minutes. The particle suspension was then centrifuged at the desired speed to assemble granular hydrogel pellets. After removing excess precursor solution in the supernatant, spheroids were gently mixed in whilst taking care to avoid bubble formation. The spheroid, particle, and interstitial matrix mixture was then transferred to cylindrical molds prepared by punching out PDMS slabs, and exposed to visible light at 20 mW.cm^-2^ for 2 minutes to crosslink the interstitial matrix. These constructs were washed thrice with culture media (Endothelial basal medium, Lonza) before culturing in culture media supplemented with 100 ng.mL^-1^ rhVEGF, 500 nM S1P, and 600 ng.mL^-1^ PMA, as previously reported.^[39]^ We confirmed that the inclusion oft he interstitial matrix did not affect particle packing and resulting interstial volume (Figure S6). The constructs were stimulated with this media for three days with media changes every day, before washing with PBS and fixation with 4% PFA. Constructs were then permeabilized with 0.1% Triton-X, blocked with 3% v/v horse serum, and incubated overnight with Alexa Fluor 647 Phalloidin at 4 °C. An upright confocal microscope (Leica TCS SP5) equipped with a 25x water immersion objective lens was used to acquire volumetric confocal stacks focusing on a single spheroid per ROI. Image processing and analysis was carried out in FIJI software.

### Subcutaneous invasion model

All animal experiments were carried out in accordance with protocols approved by the Corporal Michael J. Crescenz VA Medical Center Institutional Animal Care and Use Committee. Three male adult (2-month old) athymic rats were used for the study (Hsd:RH-Foxn1rnu, Engivo, Indianapolis, IN). To prepare for surgery, rats were anesthetized by inhalation of 2% isoflurane in oxygen carrier, and the dorsal skin prepped with alcohol and betadine. All animals received perioperative doses of cephalexin and Buprenorphine SR for antibiotic and pain coverage, respectively. A total of 200 µL of granular hydrogel sample was injected per dorsal subcutaneous site using 1 mL syringes equipped with 23G needles. A total of 6 sites were injected per animal. Sample placement was randomized with n=4 sites per group (spheres loose packing, spheres tight packing, rods loose packing, rods tight packing). After 1 week of injection, rats were euthanized, and samples harvested by separating from surrounding adipose tissue and/or fascia, and transferred to 4% PFA for overnight fixation. Subsequently, fixed samples were immersed overnight in increasingly concentrated solutions of sucrose in PBS before cryoembedding in OCT TissueTek using liquid nitrogen cooled 4-methyl butane. 35 µm thick cryosections were obtained on a cryotome and adhered onto Superfrost Plus slides. To evaluate cell infiltration, frozen cryosections were thawed to room temperature, hydrated with PBS, permeabilized with 0.1% Triton-X in PBS, blocked with 3% horse serum in PBS, and incubated with a 0.1% Hoechst solution for 15 minutes. After multiple washing steps, sections were imaged using an upright epifluorescence microscope equipped with a 10x objective lens. FIJI software was used to analyze cell infiltration. Cell density (%) was calculated as the fraction of Hoechst fluorescent area to the FITC-granular hydrogel area within multiple regions of interest (ROI). To quantify cell density at different infiltration depths within the granular hydrogel, four rectangular ROIs with a depth of 200 µm were marked using the ROI manager on FIJI software. After converting the image to 8-bit format and thresholding based on fluorescent signal, the cell density (Hoecht fluorescent area) within each ROI was quantified. To account for variability in cell density across independent samples, intensities from each of the four ROIs were summed up and cell density within each ROI was normalized to this value.

### Statistical Analysis

Data were analyzed on GraphPad Prism software. In vitro experiments were repeated at least three independent times. The Shapiro-Wilk test was conducted to determine normality of data points. To compare differences between two groups, a two-tailed student’s t-test was used. To compare differences between more than two groups, a one-way ANOVA with Tukey’s post-hoc test was used. Different levels of statistical significance were set at *p<0.05, **p<0.01, ***p<0.001, ****p<0.0001.

## Acknowledgements

This work was supported by the National Science Foundation through the Center for Engineering MechanoBiology STC (CMMI: 15-48571) and the UPenn MRSEC program (DMR-1720530), as well as through the National Institutes of Health (R01AR077362 to J.A.B.) and the German Science Foundation (QA 58/1–1 to T.H.Q.). The authors thank the Cell and Developmental Biology (CDB) Microscopy Core at the University of Pennsylvania for use of IMARIS software. The NSF Major Research Instrumentation Program (award NSF CHE1827457) and Vagelos Institute for Energy Science and Technology supported the purchase of the NMR machines used in this study. Schematics were created with Biorender.com.

## Supporting Information

**Figure S1:**
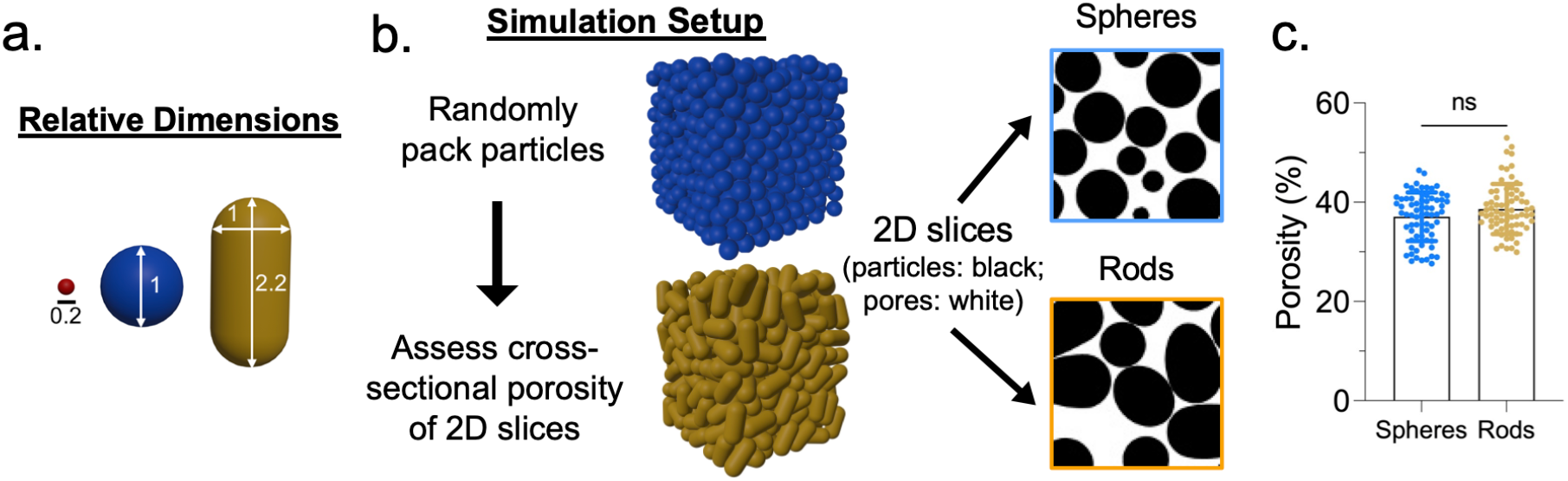
Simulation of granular assemblies using Blender. **a)** Relative dimensions of sphere (blue) and rod (gold) particles used to simulate granular assemblies (relative dimensions match those of experimental particles), as well as beads (red) used in interconnectivity simulations. **b)** Simulation of randomly packed granular assemblies from spheres (blue) and rods (gold), which are used to assess porosity by extracting 2D cross-sectional slices throughout the assembly and analyzing with image analysis (representative cross-sectional slices shown). **c)** Porosity quantification for sphere and rod-based granular assemblies. [ns = not significant]

**Figure S2:**
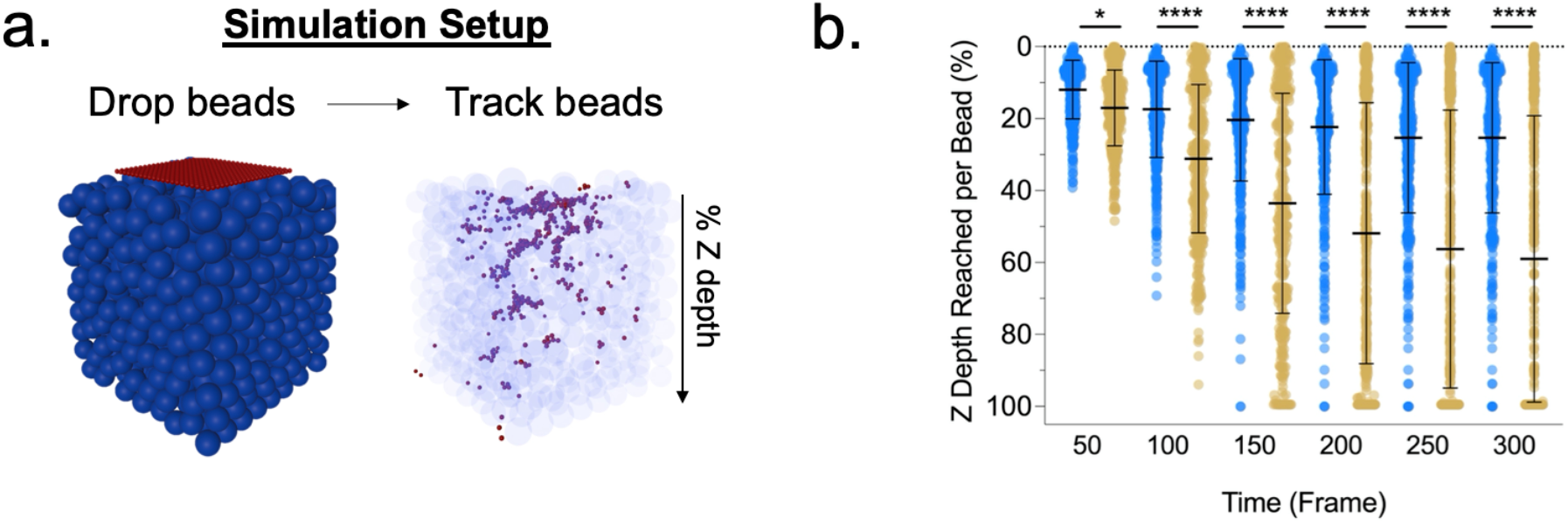
Simulation of pore interconnectivity in granular assemblies using Blender. **a)** Simulation setup consisting of dropping an array of beads and tracking bead distribution through the granular assembly over time (e.g., across frames). **b)** Distribution of Z depths reached per bead as a function of time (frame) through granular assemblies from sphere (blue) or rod (gold) particles. [*p<0.05; ****p<0.0001]

**Figure S3:**
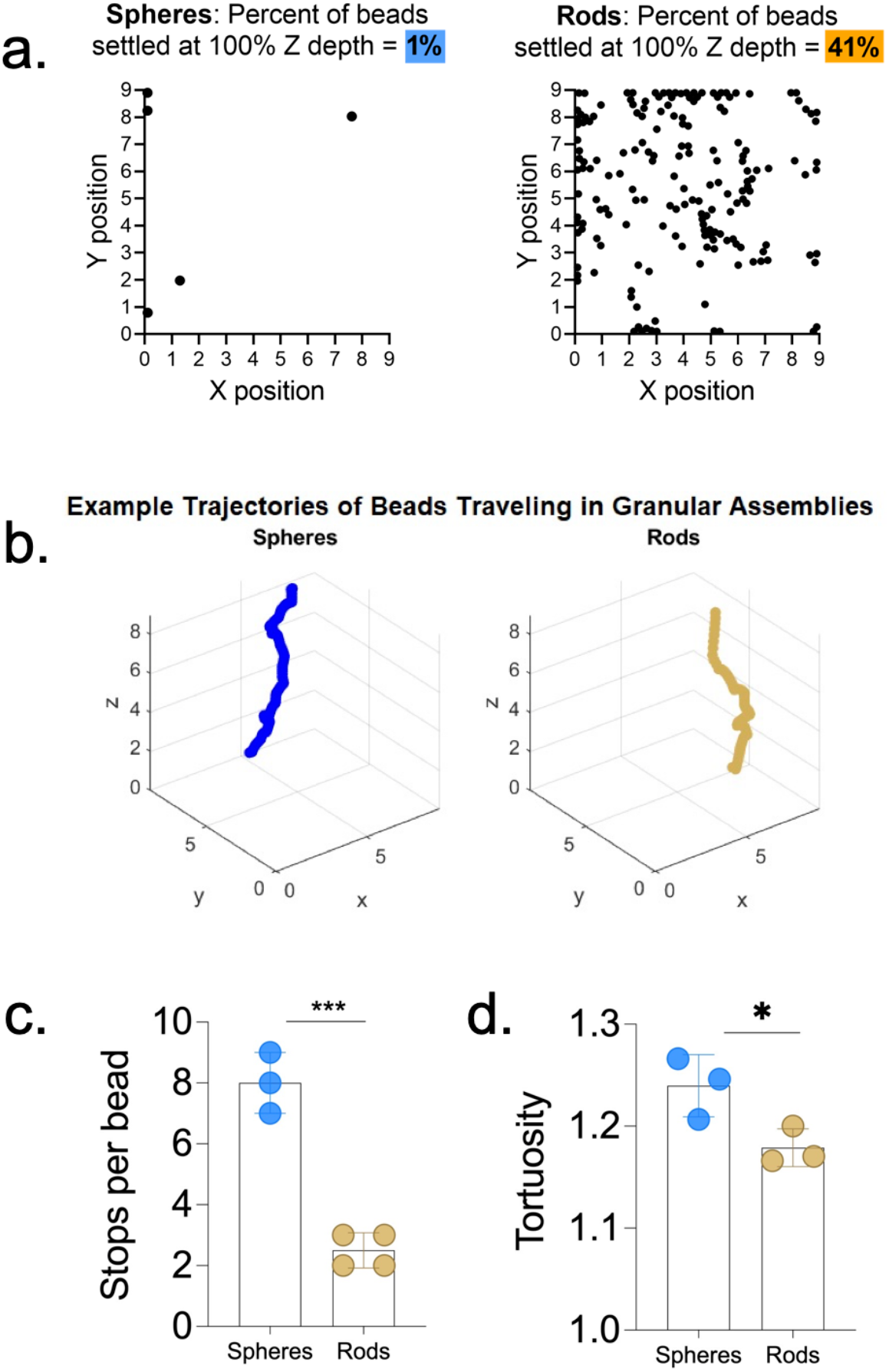
Simulation of pore interconnectivity and bead transport in granular assemblies using Blender. **a)** Distribution of beads in the XY plane that have reached 100% Z depth after equilibrium (1% of beads for spheres vs. 41% of beads for rods). **b)** Representative trajectory of a bead that traversed through the granular assembly to reach 100% Z depth. **c)** Number of times beads came to a stop while traversing through granular assemblies. **d)** Quantification of arc-chord ratio tortuosity based on bead migratory paths. [*p<0.05; ***p<0.001]

**Figure S4:**
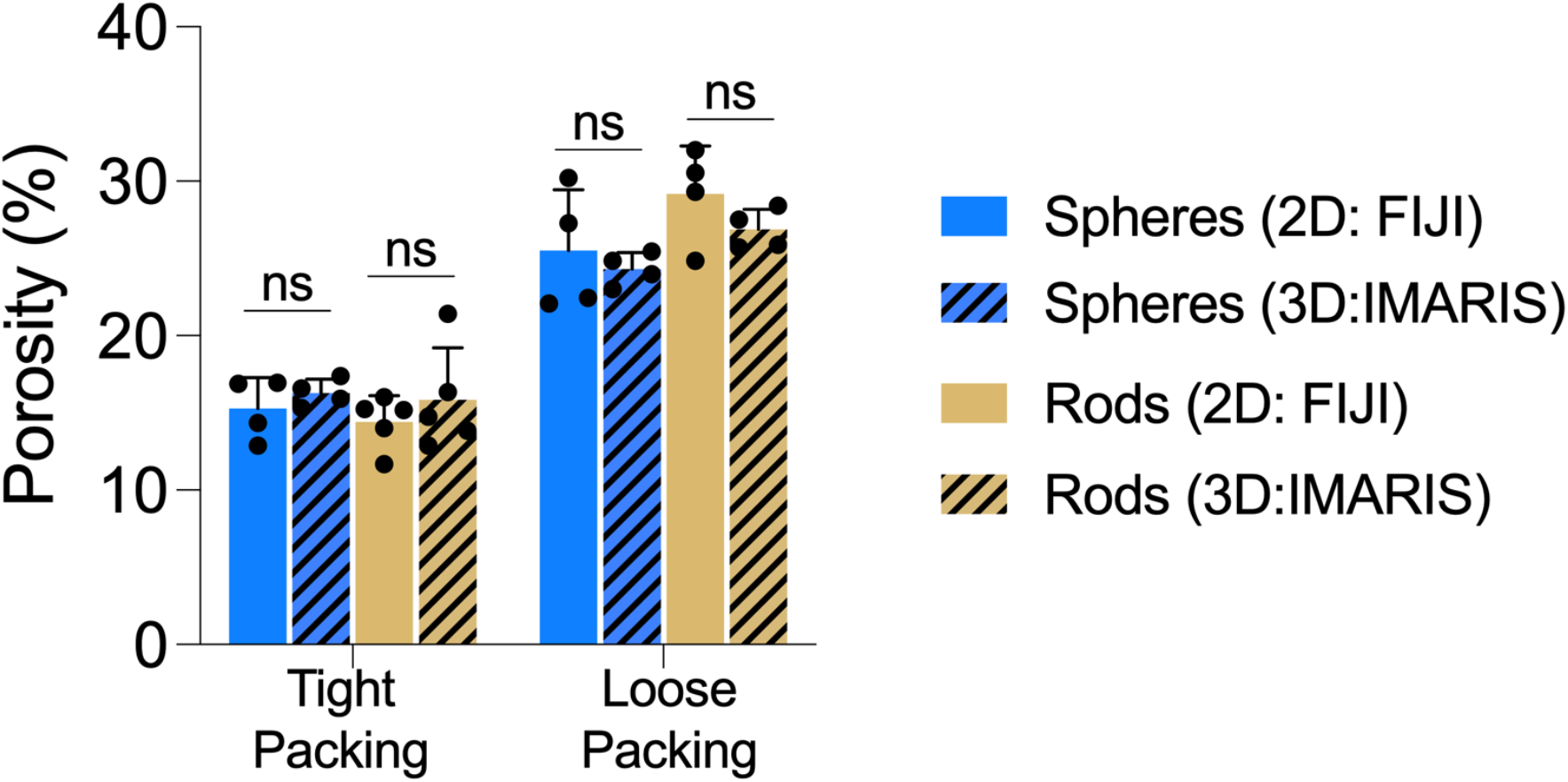
Quantified porosity of granular hydrogels. Porosity quantified using either the traditional method of thresholding 2D slices of confocal stacks (2D: FIJI) or through 3D renderings with IMARIS (3D: IMARIS). No differences are observed between the quantification methods. [ns = not significant]

**Figure S5:**
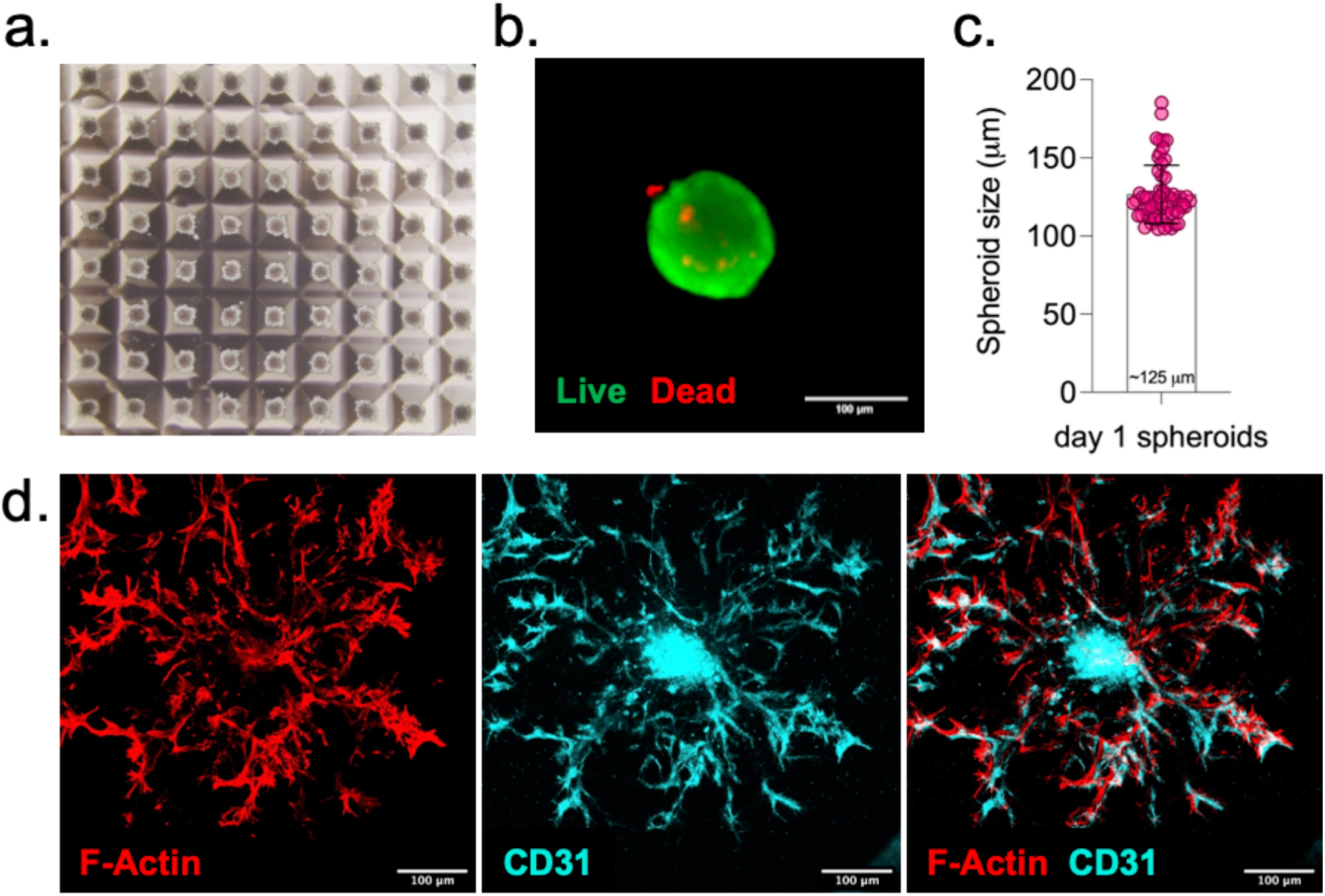
Spheroids used for endothelial sprouting assay. (a) Representative micrograph showing an array of self-assembled spheroids of HUVECs and MSCs (2:1) 24 hours after seeding on micro-templated AggreWell™ 400 well plates. (b) Live/Dead staining using Calcein AM and Propidium Iodide, respectively, on the spheroids after 24 hours. (c) Spheroids are on average ∼125 µm in diameter after 24 hours. (d) Immunofluorescent staining confirming that CD31^+^ HUVECs are located along the length of the sprouts that emerge from spheroids when embedded in a 3D degradable gel (interstitial matrix). All scale bars 100 □m.

**Figure S6:**
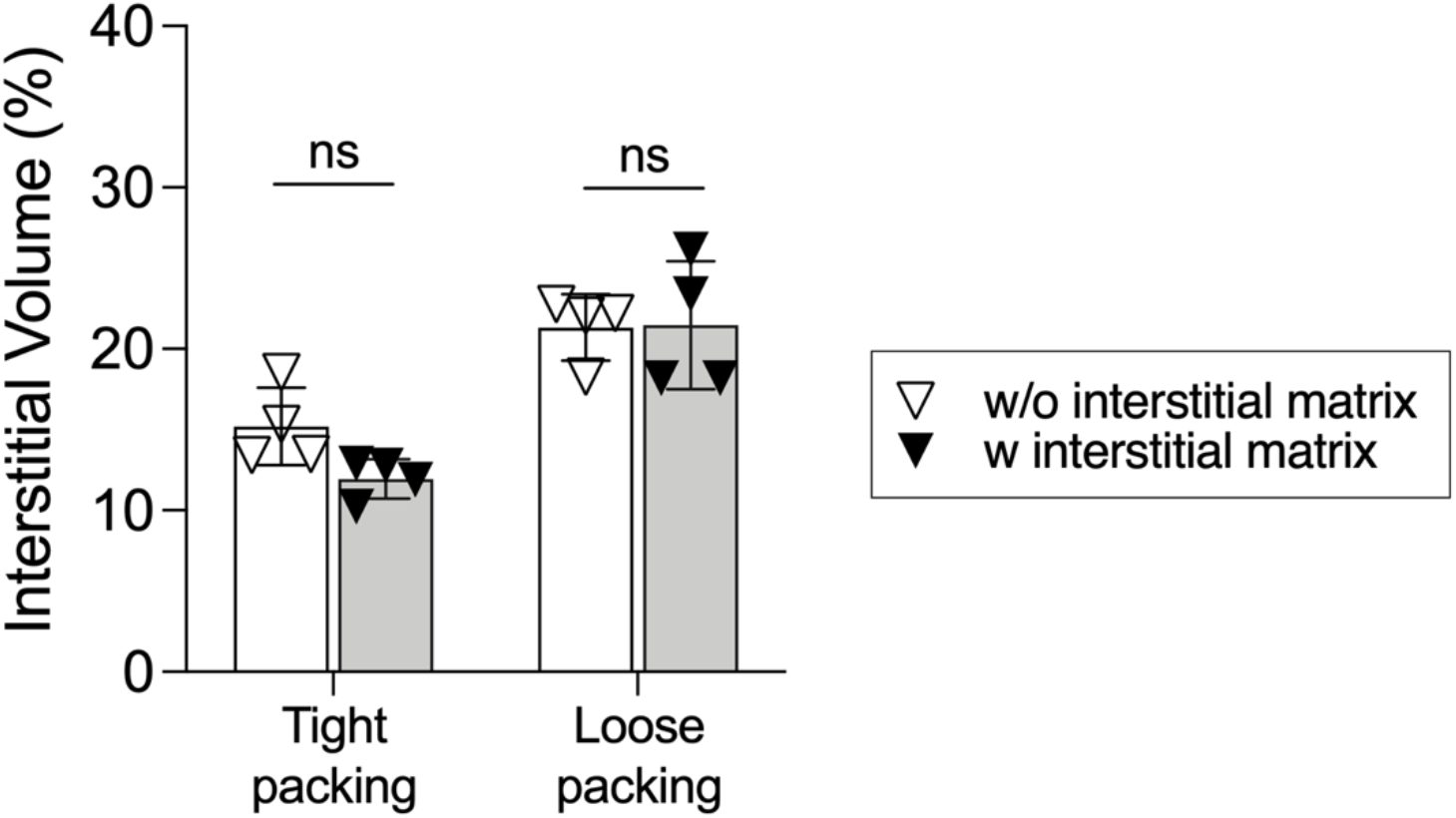
Influence of interstitial matrix on granular hydrogel interstitial volume. Quantification of granular hydrogel (from spherical particles) interstitial volume at varied particle packing densities and either with or without the inclusion of an interstitial matrix. The addition of interstitial matrix does not affect overall interstitial volume. [ns = not significant]

**Figure S7:**
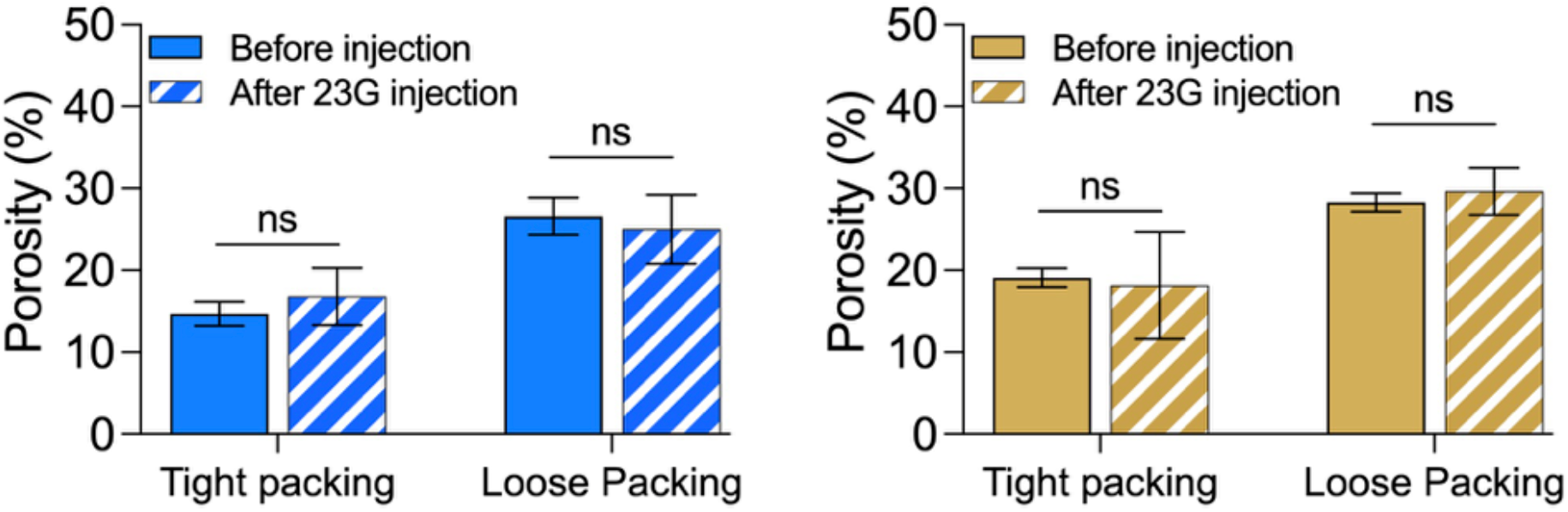
Influence of injection on granular hydrogel porosity. Quantification of granular hydrogel (from spheres (left) or rods (right)) porosity at varied packing densities and either before or after injection through 23G syringe needles. Injection does not affect overall porosity. [ns = not significant]

## Notes

### Competing Interest Statement

The authors have declared no competing interest.

